# A Watershed Algorithm GUI for Personalized fMRI-guided rTMS Target

**DOI:** 10.1101/2024.11.10.622836

**Authors:** Zi-Jian Feng, Ziyu Wei, Liquan Hong, Hongli Fang, Yu Han, Peifeng Yang, Dongsheng Lv, Yu-Feng Zang

## Abstract

Personalized repetitive transcranial magnetic stimulation (rTMS) increasingly relies on resting-state functional magnetic resonance imaging (fMRI) to select stimulation sites, yet most pipelines depend on user-defined thresholds and atlas masks, which can shift individualized targets. We propose a watershed-based approach, implemented in a graphical user interface, that performs threshold-independent segmentation of functional images to support rTMS target localization. As a proof-of-concept, we focused on Alzheimer’s disease–related circuits within the default mode network, designating the posterior cingulate cortex (PCC) as the deep effective region and the inferior parietal lobule (IPL) as the superficial stimulation target. In a cohort of 21 healthy participants, quantitative comparison with a conventional threshold-based, mask-constrained peak strategy revealed high concordance for PCC peaks but a median spatial displacement of 6.0 mm (95% CI: 0.0–12.7 mm) for IPL targets. Qualitative examples further illustrate that watershed segmentation reduces bias from neighboring functional clusters, truncation by atlas boundaries, and ambiguity among multiple local peaks. By decoupling target definition from user-chosen thresholds and packaging the method in an accessible toolbox, this framework offers a generalizable tool for individualized fMRI-guided rTMS.

## 1. Introduction

Repetitive transcranial magnetic stimulation (rTMS) is a non-invasive neuromodulation technique increasingly used for the treatment of various neuropsychiatric disorders. One of the most crucial factors influencing the efficacy of rTMS is the selection of the stimulation target (Cash et al., 2021b). In recent years, precise localization of stimulation targets based on functional magnetic resonance imaging (fMRI) is emerging as a promising approach for individualized targeting (Cash and Zalesky, 2024) (Table 1). Both functional connectivity (FC) (Feng et al., 2022; Feng et al., 2023; Siddiqi et al., 2021) and effective connectivity (Morriss et al., 2024) of resting-state fMRI (RS-fMRI) have shown effective therapeutic potential in enhancing rTMS outcomes. Additionally, task-based activation is being investigated for its ability to identify functionally relevant targets (Hong et al., 2025; Wang et al., 2023).

**Table 1.**
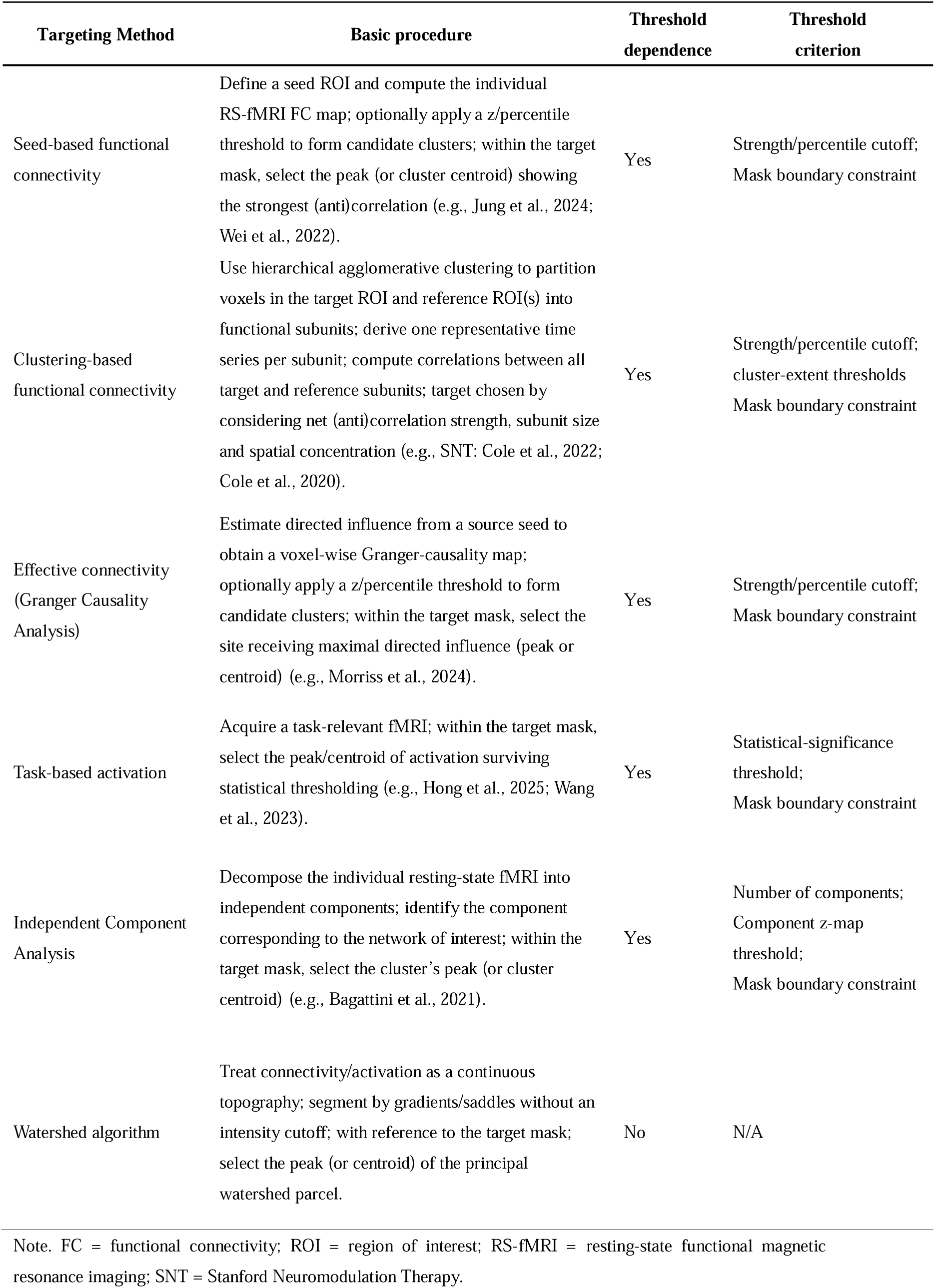
Overview of common personalized fMRI-guided TMS targeting methods.

| Targeting Method | Basic procedure | Threshold dependence | Threshold criterion |
| --- | --- | --- | --- |
| Seed-based functional connectivity | Define a seed ROI and compute the individual RS-fMRI FC map; optionally apply a z/percentile threshold to form candidate clusters; within the target mask, select the peak (or cluster centroid) showing the strongest (anti)correlation (e.g., Jung et al., 2024; Wei et al., 2022). | Yes | Strength/percentile cutoff;<br>Mask boundary constraint |
| Clustering-based functional connectivity | Use hierarchical agglomerative clustering to partition voxels in the target ROI and reference ROI(s) into functional subunits; derive one representative time series per subunit; compute correlations between all target and reference subunits; target chosen by considering net (anti)correlation strength, subunit size and spatial concentration (e.g., SNT: Cole et al., 2022; Cole et al., 2020). | Yes | Strength/percentile cutoff;<br>cluster-extent thresholds<br>Mask boundary constraint |
| Effective connectivity (Granger Causality Analysis) | Estimate directed influence from a source seed to obtain a voxel-wise Granger-causality map; optionally apply a z/percentile threshold to form candidate clusters; within the target mask, select the site receiving maximal directed influence (peak or centroid) (e.g., Morriss et al., 2024). | Yes | Strength/percentile cutoff;<br>Mask boundary constraint |
| Task-based activation | Acquire a task-relevant fMRI; within the target mask, select the peak/centroid of activation surviving statistical thresholding (e.g., Hong et al., 2025; Wang et al., 2023). | Yes | Statistical-significance threshold;<br>Mask boundary constraint |
| Independent Component Analysis | Decompose the individual resting-state fMRI into independent components; identify the component corresponding to the network of interest; within the target mask, select the cluster's peak (or cluster centroid) (e.g., Bagattini et al., 2021). | Yes | Number of components;<br>Component z-map threshold;<br>Mask boundary constraint |
| Watershed algorithm | Treat connectivity/activation as a continuous topography; segment by gradients/saddles without an intensity cutoff; with reference to the target mask; select the peak (or centroid) of the principal watershed parcel. | No | N/A |

Note. FC = functional connectivity; ROI = region of interest; RS-fMRI = resting-state functional magnetic
resonance imaging; SNT = Stanford Neuromodulation Therapy.

Previous studies have employed a framework that distinguishes a deep “effective region”, a circuit hub central to pathophysiology yet typically beyond the direct reach of figure-8 coils, from a superficial cortical “stimulation target” selected to engage that circuit via its connectivity (Feng et al., 2022; Jing et al., 2020). This general approach has been successfully applied across multiple disorders. In Alzheimer’s disease (AD), for instance, the hippocampus is frequently designated as the effective region, with the lateral parietal cortex serving as the stimulation target. High-frequency rTMS to individualized lateral parietal targets functionally connected to the hippocampus has been reported to yield short-term improvements in cognitive scores and default-mode network (DMN) connectivity (Jung et al., 2024; Wei et al., 2022). Similarly, in major depressive disorder (MDD), the most extensively studied application, the subgenual anterior cingulate cortex (sgACC) acts as the effective region, and personalized stimulation of the dorsolateral prefrontal cortex (DLPFC) based on sgACC-DLPFC FC is associated with higher response and remission rates in advanced protocols like Stanford Neuromodulation Therapy (SNT) protocol (Cole et al., 2022; Cole et al., 2020).

Despite their promise, current fMRI-guided targeting pipelines often incorporate user-defined thresholds at multiple stages. Candidate target regions are typically delineated using correlation cutoffs or percentiles, cluster-extent thresholds, and mask boundary constraints (Table 1). Small parameter changes can yield markedly different clusters and shift personalized targets (Thirion et al., 2014; Zhao et al., 2022). These issues recur across approaches such as seed-based functional connectivity (Jung et al., 2024; Wei et al., 2022), clustering-based functional connectivity (Cole et al., 2022; Cole et al., 2020), effective connectivity (Morriss et al., 2024), task-based activation (Hong et al., 2025; Wang et al., 2023) and independent component analysis (ICA) (Bagattini et al., 2021), all of which typically involve thresholding to define regions of interest (ROIs). This reliance on user-defined thresholds introduces subjectivity and complicates standardization across studies, posing challenges for reproducibility (Cash et al., 2021a; Garrison et al., 2015; Jia et al., 2021).

To ground our clinical rationale, we specify an application in AD targeting the DMN. Converging evidence identifies the posterior cingulate cortex (PCC) and precuneus as critical abnormal DMN hubs (He et al., 2015; Pan et al., 2017), a finding further supported by our recent multimodal work revealing convergent abnormalities in the dorsal precuneus in subjective cognitive decline (Li et al., 2024). We therefore designate PCC as the deep effective region and select inferior parietal lobule (IPL) as the superficial stimulation target for three reasons: 1) standard figure-8 coils cannot directly engage PCC because of depth constraints; 2) IPL is a superficial DMN hub with strong intrinsic coupling to PCC; and 3) individualized IPL stimulation has shown short-term clinical and network benefits in AD cohorts.

To mitigate threshold sensitivity and ambiguities in peak or centroid selection, we employ the watershed algorithm (Meyer, 1994), which provides a strong alternative (hereafter WSH). This algorithm conceptualizes an image as a topographic surface, where voxel intensity represents height. By simulating the progressive flooding of this surface, the algorithm identifies watershed lines that delineate catchment basins, each representing a distinct functional cluster. Crucially, this procedure is threshold-independent, the segmentation is determined by the intrinsic geometry of the connectivity distribution rather than an arbitrary cutoff. The watershed approach is therefore more robust in delineating cluster boundaries, even in the presence of multiple neighboring peaks, and it provides a principled way to define individualized ROIs for targeting without bias from threshold selection.

We implemented WSH in a user-friendly graphical interface (GUI) to create an accessible pipeline for personalized TMS target identification. Although demonstrated here for AD targeting the DMN, both the WSH algorithm and GUI are network- and diagnosis-agnostic. By allowing user-swappable effective region–stimulation target pairs, the framework provides a generalizable solution for personalized neuromodulation across diverse circuits and disorders.

## 2. Materials and Methods

### Datasets

This study utilized datasets from previous research (Yuan et al., 2018), which are freely accessible via NITRC (https://www.nitrc.org/projects/reliability). These datasets were collected with approval from the Ethics Committee of the Center for Cognition and Brain Disorders (CCBD) at Hangzhou Normal University (HZNU). RS-fMRI and 3D-T1 data were obtained from 21 healthy participants (mean age: 21.8 ± 1.8 years; 11 females), all of whom had no history of neurological, psychiatric, or neuropsychological disorders. Written informed consent was obtained from each participant prior to their inclusion in the study.

### Preprocessing of RS-fMRI

The preprocessing pipeline for RS-fMRI data included the following steps: slice timing correction, motion correction, spatial normalization to the standard Montreal Neurological Institute (MNI) space, and spatial smoothing. For identifying the stimulation target, additional preprocessing steps were performed after spatial normalization, including the regression of head motion, white matter (WM), and cerebrospinal fluid (CSF) signals, as well as the removal of linear trends. Whole-brain FC maps were Fisher-z transformed.

### Exemplar Watershed-based Implementation Demonstration

In this study, we demonstrated a pipeline for selecting rTMS targets for Alzheimer’s disease, guided by RS-fMRI FC data from healthy participants served as an illustrative case, while the pipeline is currently being applied to Alzheimer’s patients in registered studies at Chictr.org.cn (ChiCTR2400086141 and ChiCTR2400087497). In our approach, we selected the PCC as the effective region and the IPL as the stimulation target, based on their well-established roles as core hubs of the DMN (Yeo et al., 2011). The workflow of the study is illustrated in Fig. 1.

**Fig. 1.**
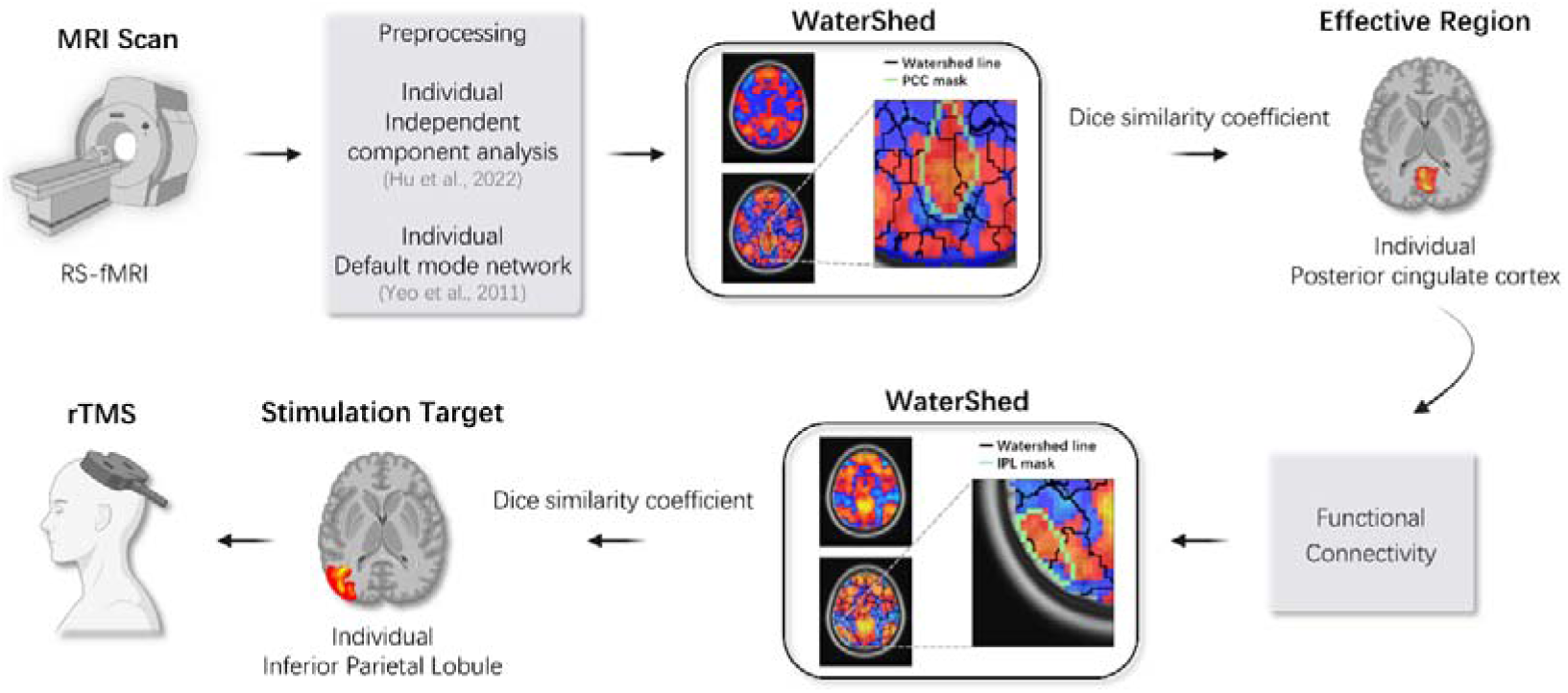
Workflow for personalized watershed-based rTMS target identification based on RS-fMRI. Following data acquisition and preprocessing, independent component analysis (ICA) identifies each individual’s default mode network (DMN). Watershed segmentation is then applied to define both the effective region and stimulation target. The effective region is determined as the peak within the cluster showing the highest Dice similarity between the PCC and individual watershed clusters within the DMN. PCC seed-based functional connectivity analysis is subsequently conducted to locate a stimulation target within the inferior parietal lobule (IPL). The final individualized stimulation target for rTMS is defined by the peak within the IPL cluster that shows the highest Dice similarity to individual watershed clusters within the DMN.

### Definition of Reference Masks

As a network prior, we used the default mode network (DMN) from Yeo et al.’s 7-network cortical parcellation (Yeo et al., 2011) as provided in the DPABI template (Yan et al., 2016). The DMN probability map was thresholded and binarized, then resampled to 3-mm isotropic MNI152 space. Connected-component labeling was applied to isolate the midline posterior component encompassing the posterior cingulate cortex and precuneus, which served as the PCC reference mask, and the left lateral-parietal component, which served as the IPL reference mask. These masks served exclusively as spatial priors, not as ground truths.

### Identification of the Effective Region (PCC)

Subject-specific spatial maps of DMN were first derived using individual ICA (Hu et al., 2022). The WSH was subsequently applied to each participant’s ICA-derived DMN component map via the GUI to parcellate the continuous functional landscape into discrete, hierarchically organized clusters. In this approach, voxel values are interpreted as terrain elevations, with lower values corresponding to valleys and higher values corresponding to peaks. The algorithm simulates a flooding process where water floods from the valleys until it encounters adjacent regions. The points at which different waters meet form “watershed line,” which effectively delineate boundaries between distinct regions, even in the presence of noise (Zhu et al., 2021). While the GUI supports visual inspection and manual selection of watershed parcels, we opted for the Dice similarity coefficient (DSC) as a more objective criterion for identifying the optimal cluster. The effective region was therefore defined as the cluster exhibiting the highest DSC with the reference PCC mask. The DSC equation is given as follows (Burunat et al., 2016):

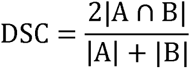

Where *A* is the set of voxels in the watershed cluster, and *B* is the set of voxels in the reference region.

The peak coordinate within this selected cluster was thus designated as the subject-specific coordinate for the effective region (i.e., PCC) and served as the seed for the subsequent connectivity analysis.

### Identification of the Stimulation Target

A seed-based FC analysis was performed using a 5 mm spherical seed region centered on the individualized PCC coordinate obtained in the previous step. This generated a whole-brain FC map for each participant. The WSH was then applied again to this connectivity map to segment it into distinct functional clusters. The stimulation target was defined as the peak coordinate of the cluster of the IPL that demonstrated the highest DSC with the reference IPL mask.

### Mask-constrained seed-based comparator

For comparison with the WSH pipeline, we implemented a seed-based FC mask-constrained peak strategy (hereafter referred to as Threshold-based, THD). Specifically, the effective region (PCC) was defined as the peak coordinate within the PCC mask on the individual’s ICA-derived DMN component map. Similarly, the stimulation target (IPL) was defined as the peak coordinate within the IPL mask on the individual’s PCC-seed FC map.

### Group-level analysis of peak selection

To directly compare the peak coordinates identified by the WSH and THD methods, we conducted a group-level analysis. For each participant and region of interest (PCC and IPL), we computed the Euclidean distance between the WSH and THD peaks. The distribution of these distances was summarized across the cohort using the median and its bias-corrected and accelerated (BCa) 95% confidence interval (10,000 resamples).

## 3. Results

Quantitative comparison demonstrated distinct spatial concordance patterns between the WSH and THD methods across regions (Fig. 2). In the PCC, the two methods showed strong agreement, with exact coordinate concordance (0 mm) in 14 of 21 participants (66.7%) and a median displacement of 0.0 mm (BCa 95% CI: 0.0–5.2 mm).

**Fig. 2.**
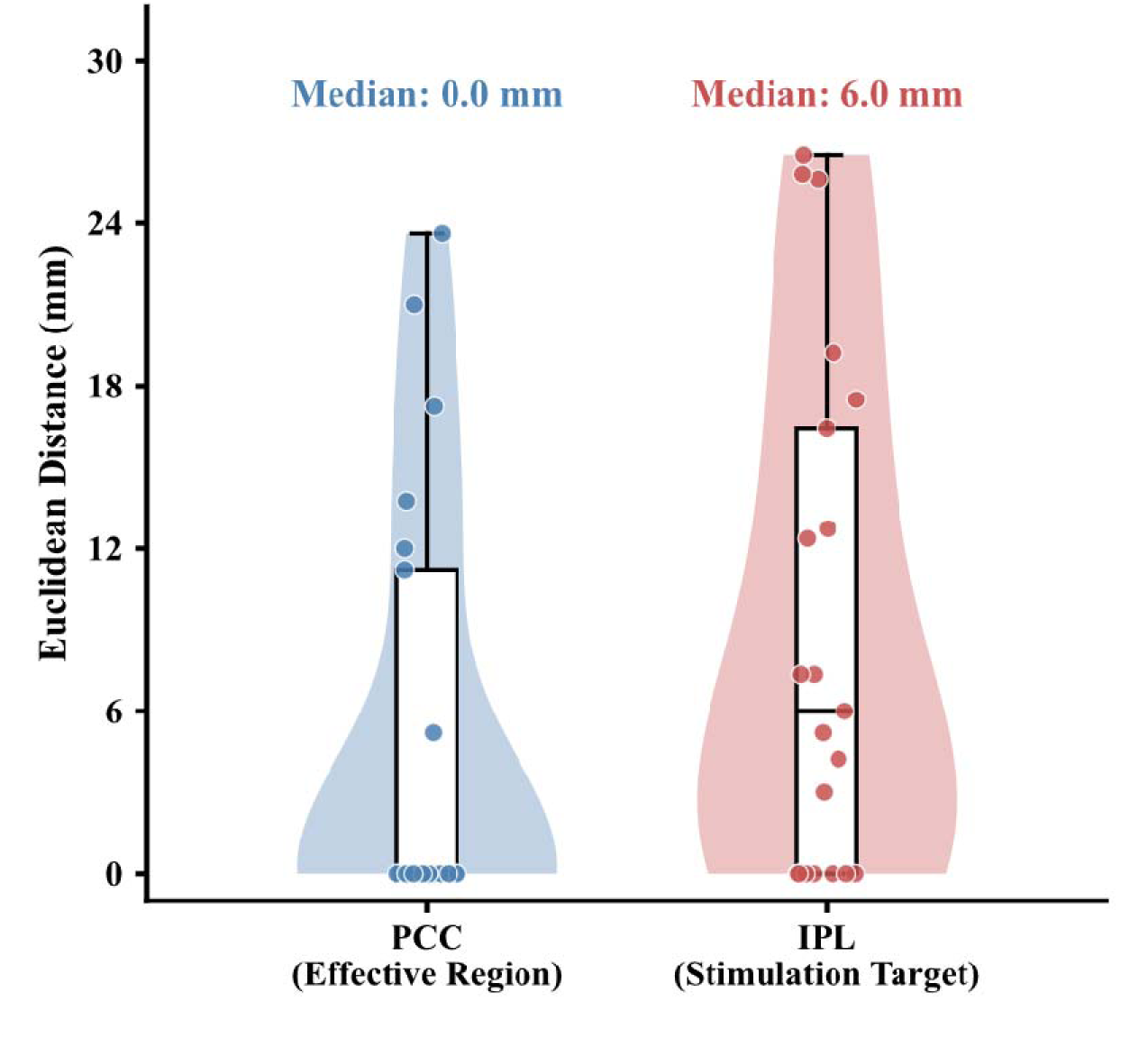
Group-level distribution of spatial offsets between watershed- and threshold-based peaks. Violin plots summarize the Euclidean distance between watershed (WSH) and threshold-based (THD) peak coordinates for the deep effective region (PCC) and the superficial stimulation target (IPL) across participants (n = 21). Individual participants are shown as dots; the central box indicates the interquartile range, the horizontal line denotes the median, and whiskers show the full range. Distances near zero indicate that WSH and THD converge on the same peak, whereas larger values reflect divergent targets. For the PCC, WSH and THD peaks were exactly concordant (0 mm) in 14 of 21 participants, yielding a median displacement of 0.0 mm. In contrast, IPL targets exhibited greater dispersion, with a median displacement of 6.0 mm and fewer exactly concordant cases (7 of 21 participants). PCC, posterior cingulate cortex; IPL, inferior parietal lobule.

In contrast, lower concordance was observed in the IPL, where different PCC seed locations introduced greater variability in the resulting functional connectivity maps. Across the full cohort, exact concordance declined to 33.3% (7/21), with a median displacement of 6.0 mm (95% CI: 0.0–12.7 mm). Importantly, this divergence was not driven solely by seed variability: even among the subset of participants (n = 14) in whom both methods used the same PCC seed, a 50% discordance rate (7/14) persisted, with a median displacement of 5.6 mm (95% CI: 0.0–12.6 mm). This indicates that the divergence stems fundamentally from how the two methods delineate the target cluster, independent of upstream seed selection. Participant-level coordinates for all subjects and spatial overlays for all discordant cases are provided in the Supplementary Materials (Tables S1–S2; Figs. S1–S2).

### Case Examples Illustrating Methodological Advantages

To qualitatively illustrate these failure modes and the corrective function of WSH, we present three representative cases. First, target selection can be biased by neighboring functional clusters when an atlas-derived mask does not align with an individual’s functional topology and inadvertently includes voxels from an adjacent parcel, importing their higher values and biasing the peak selection process within the mask toward the neighboring cluster. The THD peak localized to a different watershed parcel than the WSH-identified target. The WSH-derived parcel showed stronger spatial alignment with the reference IPL (DSC = 0.45) versus the parcel containing the THD peak (DSC = 0.16, Fig. 3). Second, truncation artifacts arise when a genuine functional cluster is cut off by a fixed mask so that the apparent peak is displaced toward the boundary (Fig. 4). Third, target ambiguity is encountered in regions with multiple local maxima, where conventional peak-picking becomes highly sensitive to minor changes in mask definition or threshold levels (Fig. 5).

**Fig. 3.**
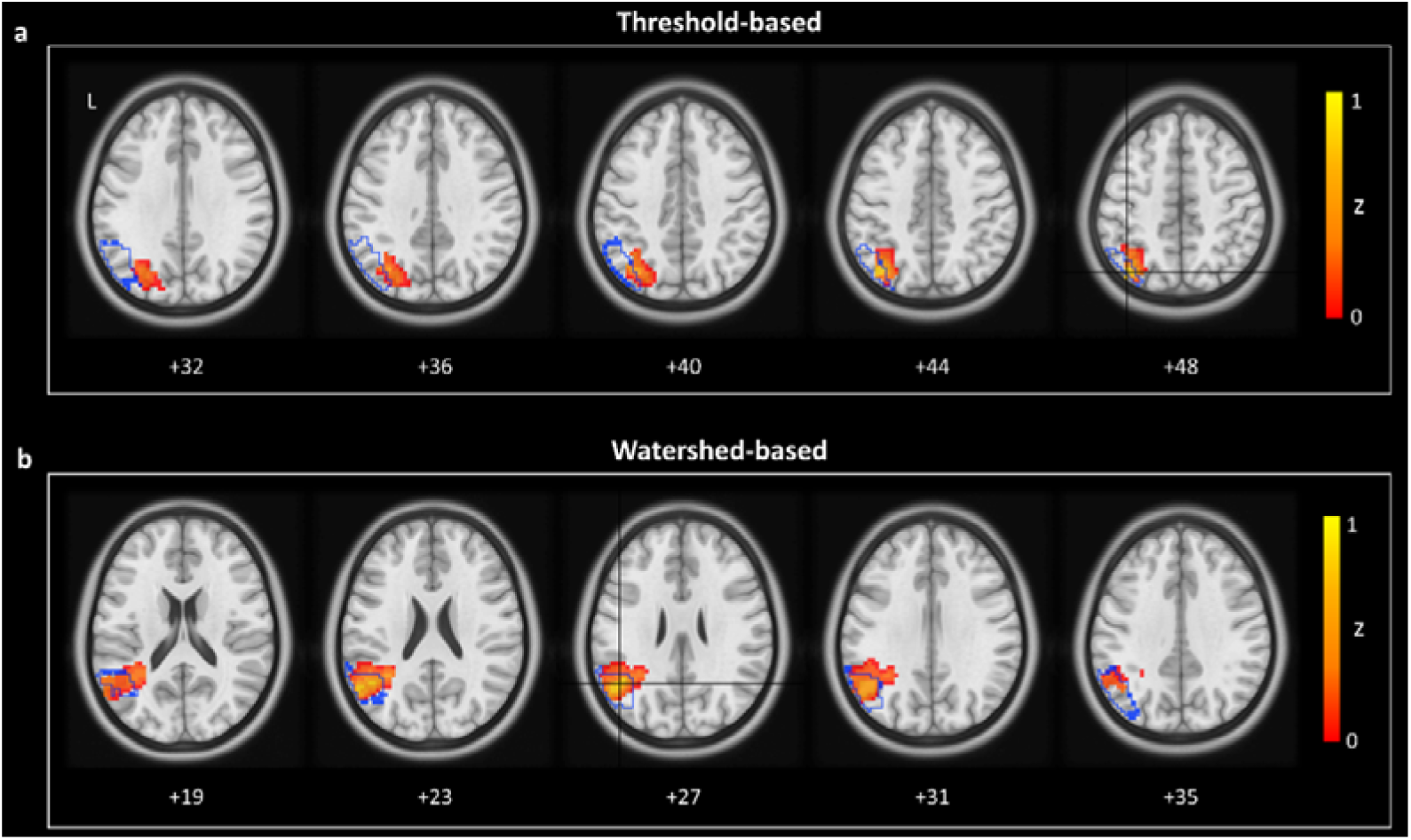
Example in which threshold-based peak selection is biased by a neighboring functional cluster, compared with watershed-based parcel selection. Axial slices show Fisher-z–transformed functional connectivity (zFC) maps from a representative participant, using the posterior cingulate cortex (PCC) as the seed. Warm colors (yellow to red) indicate positive zFC strength. The blue outline marks the atlas-defined inferior parietal lobule (IPL) mask, and the black crosshair indicates the selected stimulation target. (a) Threshold-based peak within atlas mask. The target is chosen as the within-mask voxel showing the strongest positive connectivity. In this example, the peak voxel at the crosshair is located on the edge of a larger functional cluster whose main extent lies outside the IPL mask, so the selected target is effectively driven by a neighboring cluster rather than by a parcel centered within the IPL. (b) Watershed-based parcel selection. The watershed algorithm segments the zFC map into contiguous functional parcels based on local gradients. The target is defined as the centroid of the watershed parcel that shows the greatest spatial overlap (highest Dice similarity) with the IPL mask. In this example, the resulting target is more centrally located within the overlapping parcel. All slices are shown in MNI space (z-coordinates in mm); L, left.

**Fig. 4.**
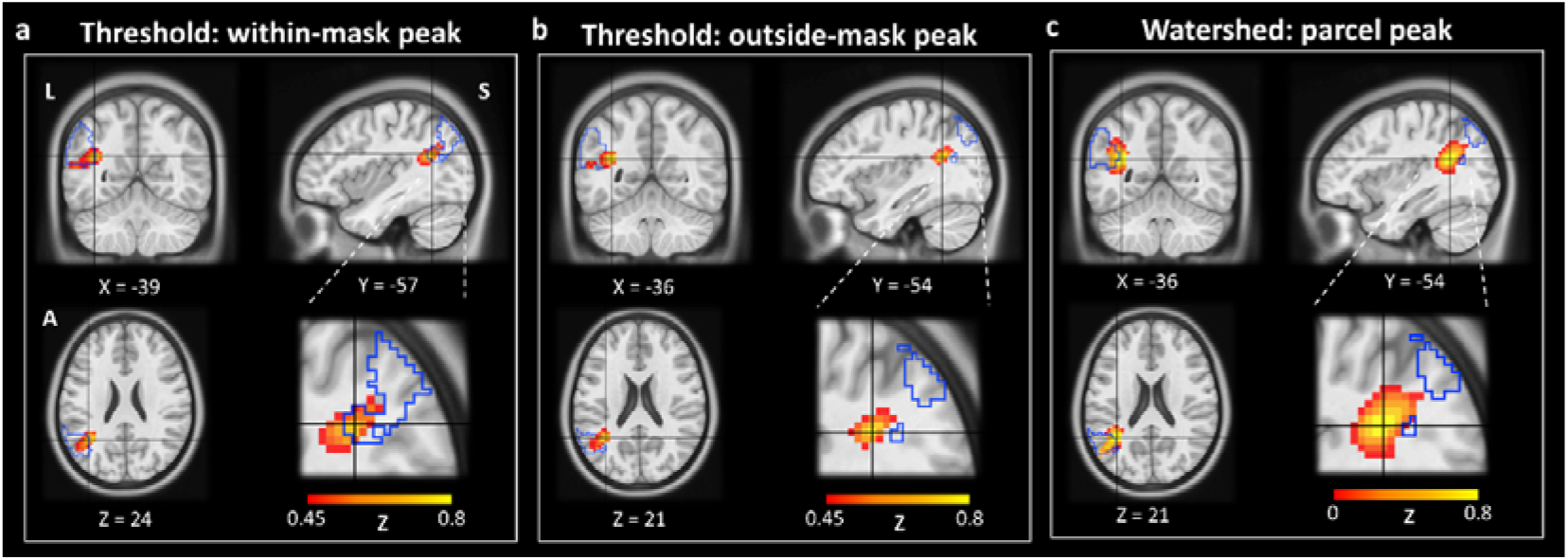
Example in which an anatomical mask truncates a functional cluster, displacing the threshold-based peak. Multi-planar views show zFC maps derived from the same PCC seed as in Fig. 3. Warm colors indicate positive zFC strength. The blue outline marks the atlas-defined inferior parietal lobule (IPL) mask, the black crosshair indicates the selected stimulation target, and dashed white lines show the zoomed-in parietal region in the insets. (a) Threshold-based peak within atlas mask. The fixed IPL mask cuts through the underlying functional cluster, truncating it at the boundary. As a result, the peak selected within the mask is displaced toward the edge of the mask, rather than reflecting the center of the cluster. (b) Threshold-based peak (unconstrained). For comparison, the peak is selected from the same zFC map without enforcing the IPL mask. The functional maximum (black cross) is located just outside the atlas boundary, indicating that the within-mask target in (a) is displaced by the mask constraint. (c) Watershed-based parcel selection. The watershed algorithm defines a functional parcel based on local gradients rather than rigid atlas edges. The resulting parcel peak closely matches the functional maximum shown in (b), reducing the displacement caused by mask truncation while remaining referenced to the IPL region. All coordinates are in MNI space (mm); A, anterior; L, left; S, superior.

**Fig. 5.**
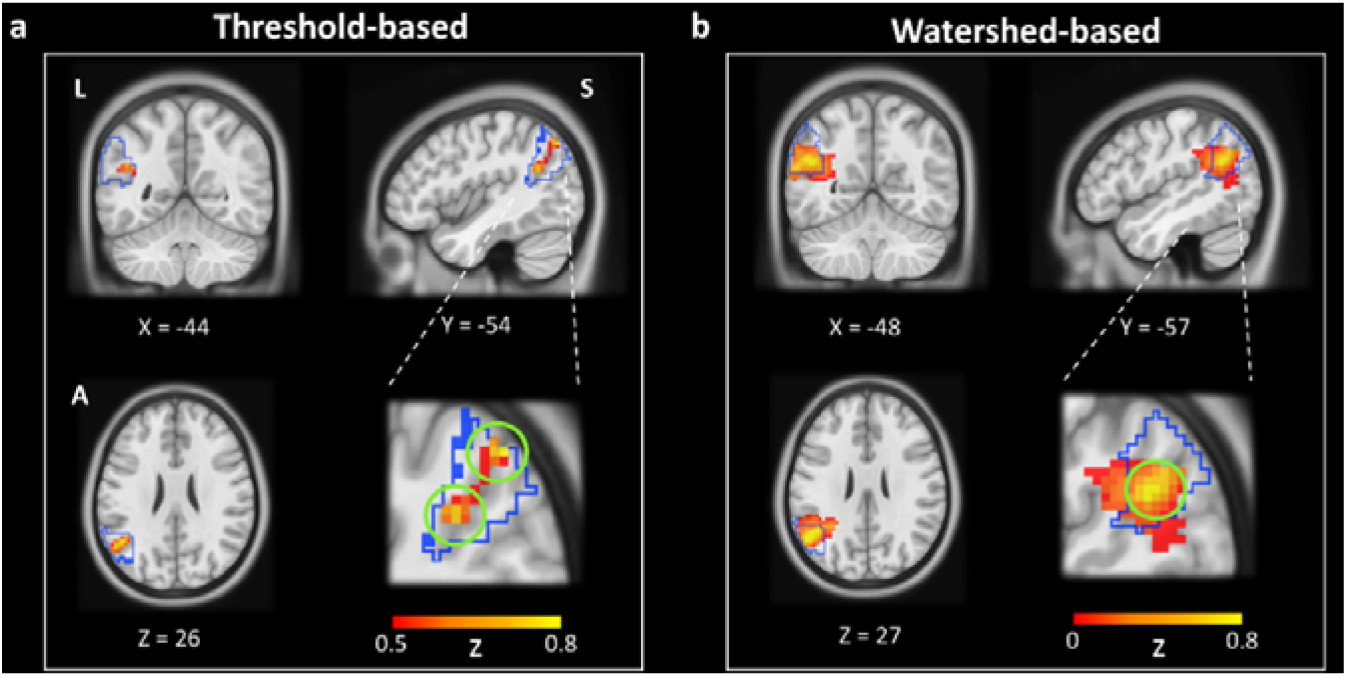
Example of target ambiguity arising from multiple local functional peaks, compared with watershed-based parcel selection. Multi-planar views show Fisher-z–transformed functional connectivity (zFC) maps derived from the same PCC seed as in Fig. 3. Warm colors indicate positive zFC strength. The blue outline delineates the atlas-derived IPL mask, green circles highlight the locations of competing local maxima (peaks) within the functional cluster, and dashed white lines indicate the zoomed parietal region in the insets. (a) Threshold-based selection. The functional landscape in this region is complex, containing several nearby local maxima (green circles) that straddle the atlas boundary. A standard threshold-based approach faces ambiguity in selecting a single “best” peak, making the target location sensitive to minor variations in signal strength or mask definition. In this case, the selected target may shift between adjacent local peaks. (b) Watershed-based selection. The watershed algorithm groups these adjacent local maxima into a single, topologically coherent parcel based on connectivity gradients. Rather than arbitrarily choosing among sub-peaks, the method defines a single target within this unified parcel (green circle), providing a more stable and centrally located stimulation site within this functional unit. All coordinates are in MNI space (mm); A, anterior; L, left; S, superior.

## 4. Discussion

This study introduces a novel method with a GUI for personalized target localization in rTMS, utilizing watershed segmentation of fMRI data. By applying the watershed algorithm to functional images and leveraging the Dice similarity coefficient, our method accurately identifies regions that optimally align with the desired target area, representing a methodological advance toward precision targeting.

In our demonstration, the PCC served as the deep effective region and the IPL as the superficial stimulation target within the DMN. In straightforward cases with a single, well-defined functional peak, the WSH and conventional THD methods converged on the same coordinate, validating the simple “peak-within-mask” approach. By contrast, when the two approaches diverge, the discrepancies are informative and trigger quantitative quality control (QC).

Our watershed framework addresses three persistent limitations of conventional, mask-constrained peak-finding: bias from adjacent functional clusters, truncation by anatomical masks, and ambiguity in local peak selection (Figs. 3–5). Although ad hoc visual review is sometimes used to mitigate these issues, it is time-consuming and adds subjectivity and inter-rater variability. These issues proved frequent in practice, affecting 33% (7/21) of PCC and 50% (7/14) of IPL targets in our cohort. In such challenging cases, our WSH method consistently established gradient-defined parcel boundaries, directly circumventing artifacts caused by cluster adjacency and mask truncation to provide a stable, threshold-independent parcellation. This approach identified the principal functional unit, even in complex cases, typically yielding parcels with improved template overlap and more topologically central peaks.

The generalizability of the watershed approach extends beyond the DMN to other circuits, such as the sgACC-DLPFC network in MDD. Although the full implementation details of advanced protocols such as SNT are not publicly disclosed (Cole et al., 2022; Cole et al., 2020), they are generally characterized by two methodological constraints: reliance on thresholded steps for subunit definition and confinement to predefined anatomical or network masks. By contrast, WSH generates candidate subunits directly from connectivity gradients, reducing threshold sensitivity and providing objective QC metrics (e.g., template Dice) without altering the underlying clinical rationale.

Furthermore, the proposed watershed framework offers a modality-agnostic solution for functional localization. While we have demonstrated its utility with RS-fMRI, the underlying principles are directly transferable to task-based fMRI. This capability is critical for obtaining reliable functional targets in clinical conditions like stroke, where compensatory plasticity and cortical reorganization can profoundly shift activation peaks and alter network topography (Cramer et al., 2001), rendering conventional template-based localization inadequate. The framework’s ability to derive targets from either resting-state or task-based data provides the flexibility needed to map these reorganized circuits faithfully.

To enhance clinical accessibility, all watershed computation steps were integrated into the GUI of the RESTplus toolkit (Jia et al., 2019). We developed a GUI pipeline that enables users without specialized coding or neuroimaging expertise to apply this method. The GUI facilitates one-click watershed segmentation by allowing users to input preprocessed individual fMRI data and specify the target region template (Fig. S3). Additionally, it provides the watershed cluster with the highest Dice coefficient and a complete list of Dice coefficients for all clusters. This streamlined, user-friendly interface significantly reduces the complexity of personalized target identification for rTMS, promoting broader adoption in clinical settings.

However, it is worth noting that the direct intervention depth of TMS is limited. Consequently, peaks located in deeper cortical regions may not be directly accessible with standard figure-8 coils. This constraint is shared across personalized fMRI-guided approaches, including the present framework (e.g., Fig. 4). Therefore, final target selection in practice requires investigators to balance optimal network engagement against practical considerations of anatomical accessibility and patient tolerability.

In summary, we have developed a watershed-based approach for guiding personalized rTMS target localization and designed a corresponding GUI toolbox. This GUI-driven pipeline lowers the barrier for clinical implementation by enabling clinicians without specialized neuroimaging expertise to effectively apply individualized neuromodulation strategies. Our method sets the stage for broader adoption of personalized rTMS interventions, enhancing accessibility and allowing for more tailored treatment approaches.

## Acknowledgments

The authors have no acknowledgments to report.

## 7. Data availability

All resources required to reproduce the findings of this study, including the Watershed GUI, source code, ROI masks, and processing parameters, are publicly available at (https://gitlab.gwdg.de/brain-networks-mpi-cbs/papers/Watershed-Algorithm-GUI). The original dataset analyzed in this study is hosted separately at NITRC (https://www.nitrc.org/projects/reliability/). Please cite this article and the repository URL when reusing these materials.

