## Supplementary material for "A Watershed Algorithm GUI for Personalized fMRI-guided rTMS Target": Figs. S1-S2; Fig. S3

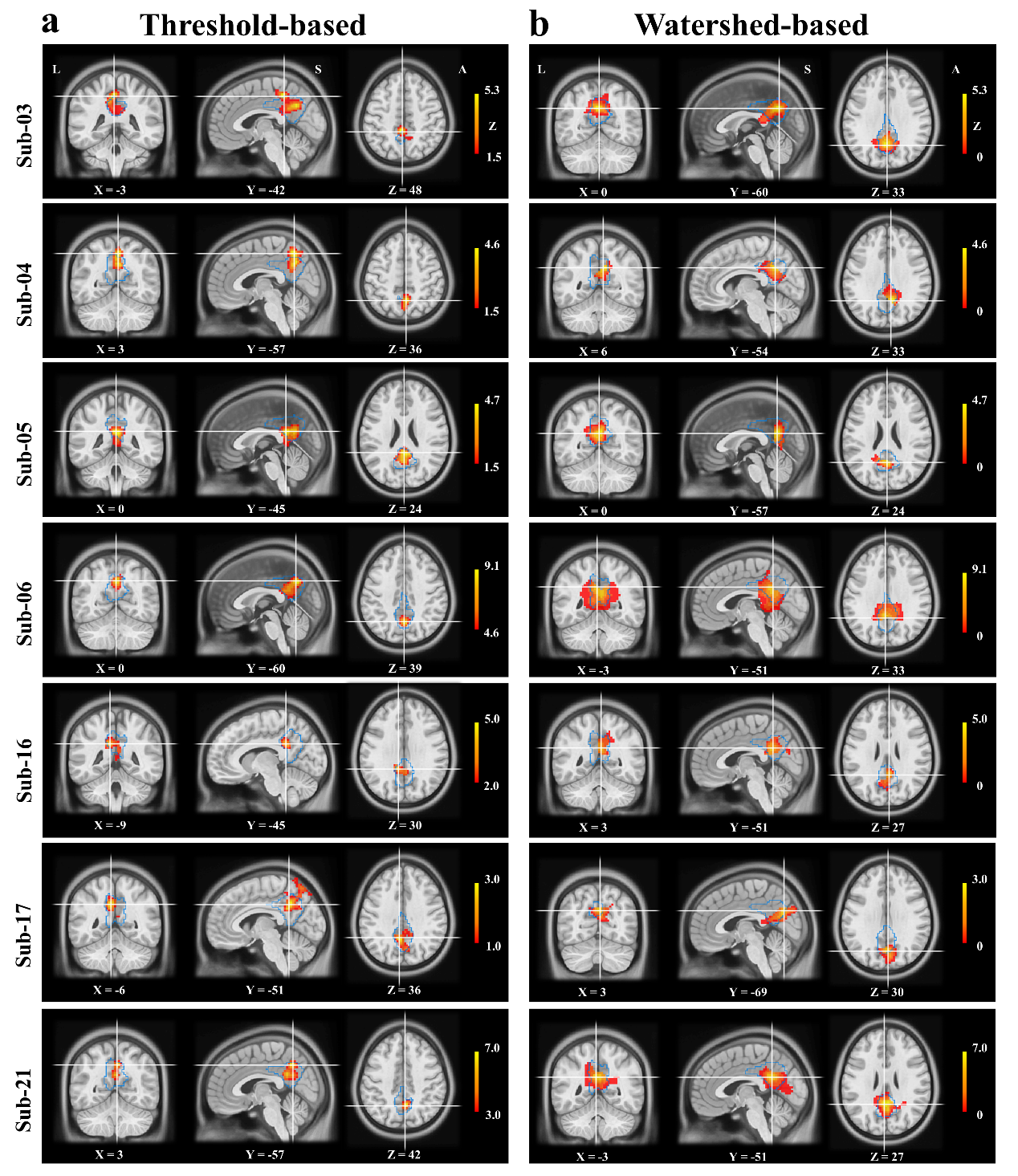

**Fig. S1. Differences in effective regions between watershed- and threshold-based approaches using subject-specific DMN across individuals.**

For each participant (rows), maps display the subject-specific ICA component corresponding to the default mode network (DMN); The blue outline marks the atlas-derived posterior cingulate cortex (PCC). (a) Threshold-based: effective regions are defined by the within-mask local maximum of a suprathreshold portion of the DMN component. (b) Watershed-based: a gradient-defined, subject-specific parcellation assigns a single coherent parcel to this region and places the effective region at the parcel peak of the parcel with the highest Dice similarity to the reference PCC mask. All coordinates are in MNI space (z-coordinates in mm); the color bars indicate zFC values; peaks are marked by a black cross; ICA, independent component analysis A, Anterior; L, Left; S, Superior.

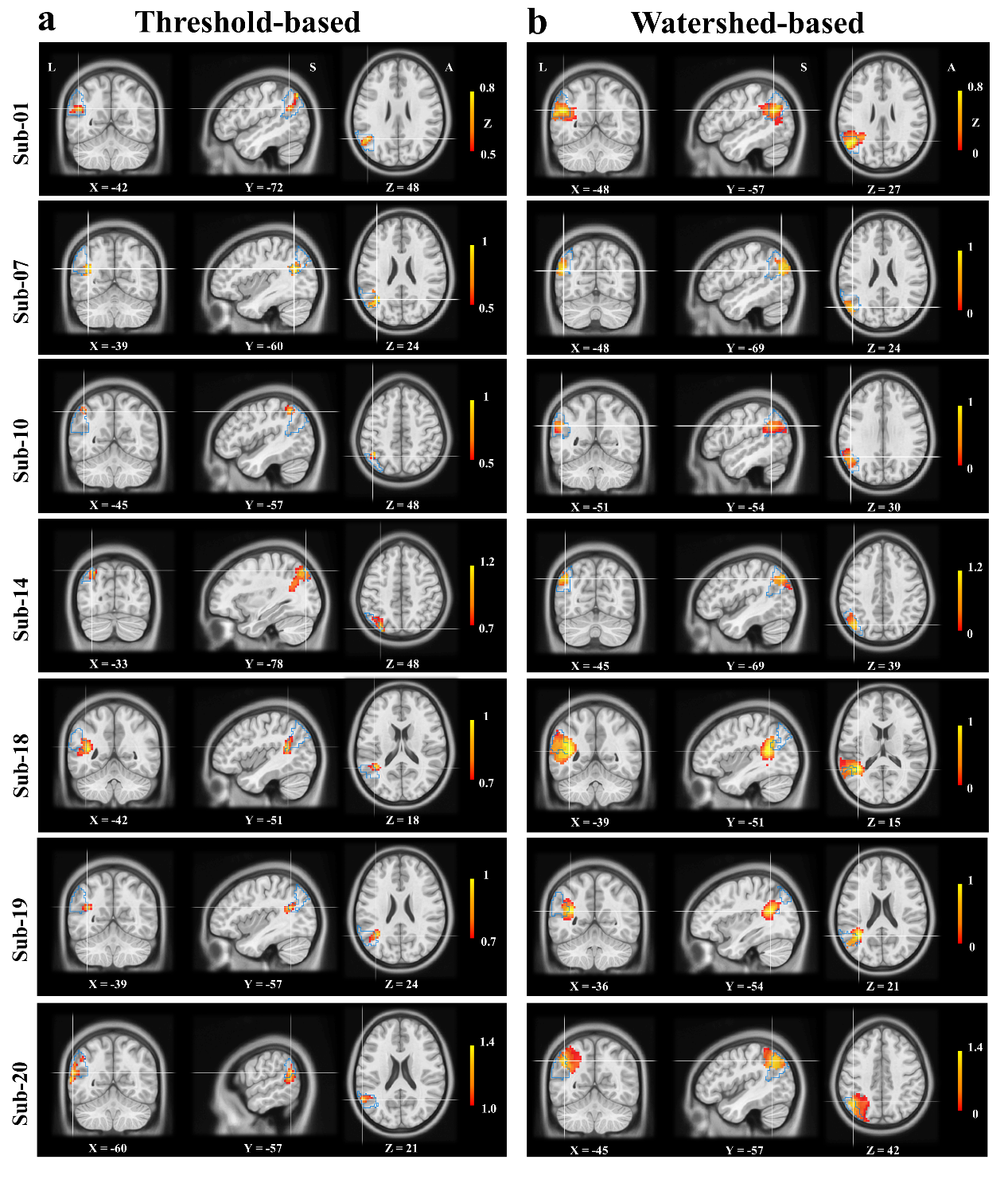

**Fig. S2. Differences in stimulation targets between watershed- and threshold-based approaches using the same PCC seed across individuals.**

Maps show Fisher-z–transformed functional connectivity (zFC) from the same posterior cingulate cortex (PCC) seed. The blue outline marks the atlas-derived inferior parietal lobule (IPL). (a) Threshold-based: the target is the within-mask local maximum from a suprathreshold cluster. (b) Watershed-based: a gradient-defined, subject-specific parcellation assigns a single parcel; the target is the parcel peak of the parcel with the highest Dice similarity to the reference IPL mask. All coordinates are in MNI space (z-coordinates in mm); the color bars indicate zFC values; peaks are marked by a black cross; A, Anterior; L, Left; S, Superior.

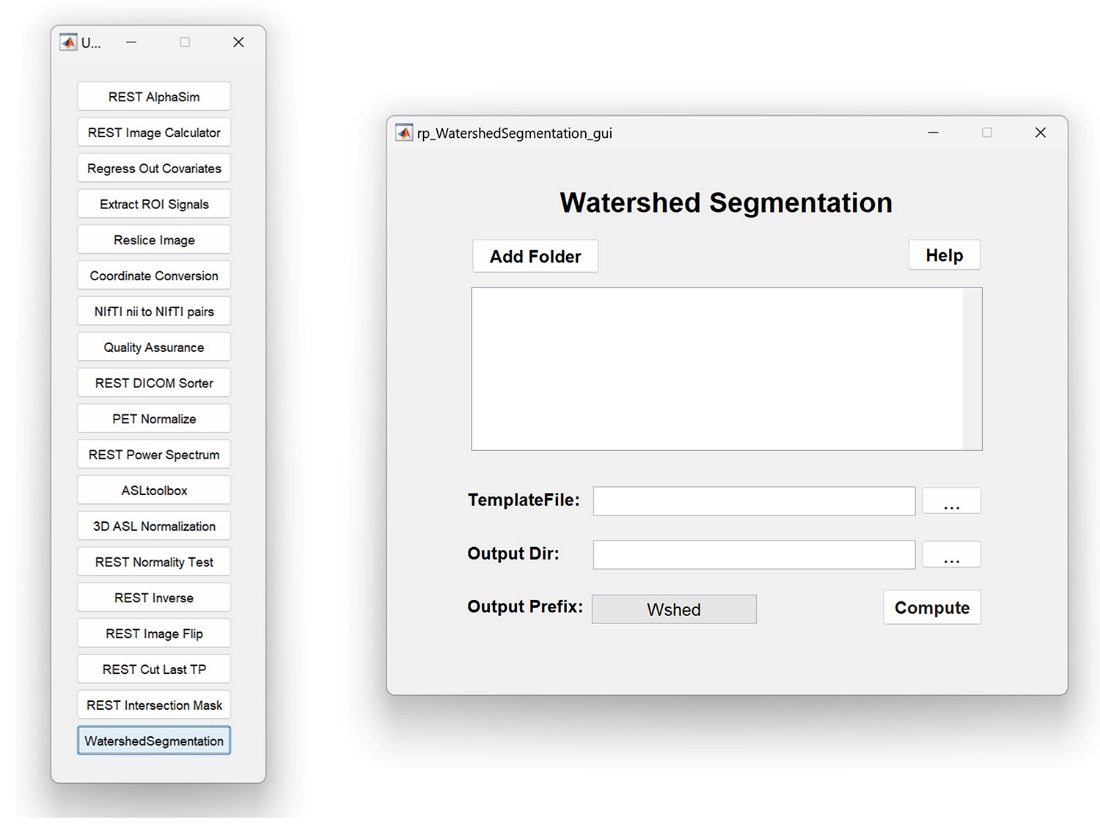

**Fig. S3. The user interface of the watershed segmentation module within the RESTplus toolkit.**

This module facilitates the segmentation process by allowing users to load folders containing fMRI data and specify template files and output directories. Key features include buttons to add folders, select template files, and set output parameters. The "Compute" button initiates the segmentation process, and users can customize the output file prefix. The interface is designed to simplify the process for users without specialized programming skills, making it accessible for clinical and research applications involving personalized fMRI-guided target identification.

Table S1. Participant-level effective region coordinates for all subjects

| **Sub ID** | **THD-PCC** | | |  | **WSH-PCC** | | | **ED** |
| --- | --- | --- | --- | --- | --- | --- | --- | --- |
|  | X | Y | Z |  | X | Y | Z |  |
| Sub 01 | -3 | -48 | 33 |  | -3 | -48 | 33 | 0 |
| Sub 02 | 9 | -57 | 42 |  | 9 | -57 | 42 | 0 |
| Sub 03 | -3 | -42 | 48 |  | 0 | -60 | 33 | 23.62 |
| Sub 04 | 3 | -57 | 36 |  | 6 | -54 | 33 | 5.20 |
| Sub 05 | 0 | -45 | 24 |  | 0 | -57 | 24 | 12.00 |
| Sub 06 | 0 | -60 | 39 |  | -3 | -51 | 33 | 11.22 |
| Sub 07 | 3 | -60 | 33 |  | 3 | -60 | 33 | 0 |
| Sub 08 | 0 | -48 | 18 |  | 0 | -48 | 18 | 0 |
| Sub 09 | -12 | -45 | 33 |  | -12 | -45 | 33 | 0 |
| Sub 10 | -9 | -54 | 39 |  | -9 | -54 | 39 | 0 |
| Sub 11 | -6 | -45 | 30 |  | -6 | -45 | 30 | 0 |
| Sub 12 | 0 | -60 | 36 |  | 0 | -60 | 36 | 0 |
| Sub 13 | -9 | -51 | 33 |  | -9 | -51 | 33 | 0 |
| Sub 14 | 0 | -48 | 21 |  | 0 | -48 | 21 | 0 |
| Sub 15 | -9 | -39 | 36 |  | -9 | -39 | 36 | 0 |
| Sub 16 | -9 | -45 | 30 |  | 3 | -51 | 27 | 13.75 |
| Sub 17 | -6 | -51 | 36 |  | 3 | -69 | 30 | 21.00 |
| Sub 18 | 0 | -66 | 24 |  | 0 | -66 | 24 | 0 |
| Sub 19 | 3 | -69 | 30 |  | 3 | -69 | 30 | 0 |
| Sub 20 | -3 | -45 | 36 |  | -3 | -45 | 36 | 0 |
| Sub 21 | 3 | -57 | 42 |  | -3 | -51 | 27 | 17.23 |

Note. THD = Threshold-based; WSH = watershed-based; PCC = posterior cingulate cortex; ED = Euclidean distance.

Table S2. Participant-level stimulation target coordinates for all subjects

| **Sub ID** | **THD-IPL** | | |  | **WSH-IPL** | | | **ED** |
| --- | --- | --- | --- | --- | --- | --- | --- | --- |
|  | X | Y | Z |  | X | Y | Z |  |
| Sub 01 | -42 | -72 | 48 |  | -48 | -57 | 27 | 26.50 |
| Sub 02 | -45 | -60 | 27 |  | -45 | -60 | 27 | 0 |
| Sub 03 | -51 | -48 | 33 |  | -42 | -72 | 33 | 25.63 |
| Sub 04 | -60 | -60 | 24 |  | -57 | -60 | 24 | 3.00 |
| Sub 05 | -39 | -75 | 27 |  | -33 | -75 | 27 | 6.00 |
| Sub 06 | -45 | -57 | 36 |  | -51 | -66 | 30 | 12.37 |
| Sub 07 | -39 | -60 | 24 |  | -48 | -69 | 24 | 12.73 |
| Sub 08 | -42 | -75 | 36 |  | -42 | -75 | 36 | 0 |
| Sub 09 | -51 | -57 | 33 |  | -51 | -57 | 33 | 0 |
| Sub 10 | -45 | -57 | 48 |  | -51 | -54 | 30 | 19.21 |
| Sub 11 | -48 | -72 | 30 |  | -48 | -72 | 30 | 0 |
| Sub 12 | -45 | -72 | 33 |  | -45 | -72 | 33 | 0 |
| Sub 13 | -57 | -66 | 27 |  | -57 | -66 | 27 | 0 |
| Sub 14 | -33 | -78 | 48 |  | -45 | -69 | 39 | 17.49 |
| Sub 15 | -54 | -60 | 33 |  | -54 | -60 | 33 | 0 |
| Sub 16 | -45 | -60 | 33 |  | -48 | -66 | 36 | 7.35 |
| Sub 17 | -39 | -66 | 36 |  | -42 | -69 | 42 | 7.35 |
| Sub 18 | -42 | -51 | 18 |  | -39 | -51 | 15 | 4.24 |
| Sub 19 | -39 | -57 | 24 |  | -36 | -54 | 21 | 5.20 |
| Sub 20 | -60 | -57 | 21 |  | -45 | -57 | 42 | 25.81 |
| Sub 21 | -39 | -63 | 39 |  | -42 | -57 | 24 | 16.43 |

Note. THD = Threshold-based; WSH = watershed-based; IPL = inferior parietal lobule; ED = Euclidean distance.
